# Deep autoencoder-based behavioral pattern recognition outperforms standard statistical methods in high-dimensional zebrafish studies

**DOI:** 10.1101/2023.09.13.557544

**Authors:** Adrian J. Green, Lisa Truong, Preethi Thunga, Connor Leong, Melody Hancock, Robyn L. Tanguay, David M. Reif

**Author notes:** Corresponding Author: E-mail Address (AJG).

## Abstract

Zebrafish have become an essential tool in screening for developmental neurotoxic chemicals and their molecular targets. The success of zebrafish as a screening model is partially due to their physical characteristics including their relatively simple nervous system, rapid development, experimental tractability, and genetic diversity combined with technical advantages that allow for the generation of large amounts of high-dimensional behavioral data. These data are complex and require advanced machine learning and statistical techniques to comprehensively analyze and capture spatiotemporal responses. To accomplish this goal, we have trained semi-supervised deep autoencoders using behavior data from unexposed larval zebrafish to extract quintessential “normal” behavior. Following training, our network was evaluated using data from larvae shown to have significant changes in behavior (using a traditional statistical framework) following exposure to toxicants that include nanomaterials, aromatics, per- and polyfluoroalkyl substances (PFAS), and other environmental contaminants. Further, our model identified new chemicals (Perfluoro-n-octadecanoic acid, 8-Chloroperfluorooctylphosphonic acid, and Nonafluoropentanamide) as capable of inducing abnormal behavior at multiple chemical-concentrations pairs not captured using distance moved alone. Leveraging this deep learning model will allow for better characterization of the different exposure-induced behavioral phenotypes, facilitate improved genetic and neurobehavioral analysis in mechanistic determination studies and provide a robust framework for analyzing complex behaviors found in higher-order model systems.

**Author Summary:** We demonstrate that a deep autoencoder using raw behavioral tracking data from zebrafish toxicity screens outperforms conventional statistical methods, resulting in a comprehensive evaluation of behavioral data. Our models can accurately distinguish between normal and abnormal behavior with near-complete overlap with existing statistical approaches, with many chemicals detectable at lower concentrations than with conventional statistical tests; this is a crucial finding for the protection of public health. Our deep learning models enable the identification of new substances capable of inducing aberrant behavior, and we generated new data to demonstrate the reproducibility of these results. Thus, neurodevelopmentally active chemicals identified by our deep autoencoder models may represent previously undetectable signals of subtle individual response differences. Our method elegantly accounts for the high degree of behavioral variability associated with the genetic diversity found in a highly outbred population, as is typical for zebrafish research, thereby making it applicable to multiple laboratories. Utilizing the vast quantities of control data generated during high-throughput screening is one of the most innovative aspects of this study and to our knowledge is the first study to explicitly develop a deep autoencoder model for anomaly detection in large-scale toxicological behavior studies.

## Introduction

Significant progress continues to be made in our understanding of neurodevelopmental disorders such as autism spectrum disorder, attention deficit hyperactivity disorder (ADHD), developmental delay, learning disabilities, and other neurodevelopmental problems. As incidences continue to rise globally and affect 10-15% of all births, more work must be done to improve our understanding of these disorders (Boyle et al., 2011; *Neurodevelopmental Diseases*, 2021; US EPA, 2015b). Meta-analyses suggest strong and consistent epidemiological evidence that the developing nervous system is particularly vulnerable to low-level exposure to widespread environmental contaminants, as the anatomical and functional architecture of the human brain is mainly determined by developmental transcriptional processes during the prenatal period (Grandjean & Landrigan, 2014; Green & Planchart, 2018; Miller et al., 2014; Rock & Patisaul, 2018; US EPA, 2015b). Therefore, identifying associations between developmental exposures and neurological effects is a core objective to improve public health by informing disease and disability prevention (*A Blueprint for Brain Development*, 2014; *Neurodevelopmental Diseases*, 2021).

As the number of environmental contaminants grows to nearly one million, comprehensive data on the neurodevelopmental toxicity of these contaminants remain sparse or nonexistent (Krewski et al., 2020; US EPA, 2015a, 2015b; Wambaugh et al., 2013). In response, high-throughput screening (HTS) assays have been developed to expedite chemical toxicity testing using *in vitro* and *in vivo* systems (Judson et al., 2010; Richard et al., 2016; Truong et al., 2014). However, *in vitro* cell and cell-free assays cannot fully capture systemic organismal responses in terms of anatomy, physiology, or behavior (Thomas et al., 2012). Zebrafish (*Danio rerio*) have emerged as an ideal model for studying low-level chemical exposure because of their high fecundity, rapid development, genetic tractability, and amenability to high-throughput data generation (Bugel et al., 2014; Planchart et al., 2018; Truong et al., 2014). The zebrafish brain’s structural organization, cellular morphology, and neurotransmitter systems are very similar to other vertebrates, including chickens, rats, and humans (Horzmann & Freeman, 2016; Kalueff et al., 2014; Lowery & Sive, 2004; Tropepe & Sive, 2003). Furthermore, zebrafish have behavioral patterns highly similar to mammals, and genetic homologs for 70% of human genes and 82% of human disease genes, making them a powerful tool for revealing the neuronal developmental pathways underlying behavior (Basnet et al., 2019; Howe et al., 2013; Postlethwait et al., 1998).

Zebrafish larvae show mature swimming patterns following swim bladder development at four to five days post-fertilization (dpf), which can be assessed using various locomotor behavioral assays (Hernandez et al., 2018; Tegelenbosch et al., 2012). One of these assays, the larval photomotor response (LPR), utilizes a sudden transition from light to dark, eliciting a stereotyped large-angle O-bend, followed by several minutes of increased movement, which gradually reduces (Burgess & Granato, 2007c; Emran et al., 2008). Exposure to toxicants has been shown to alter this stereotypical behavioral response (Basnet et al., 2019; Truong et al., 2016). Current HTS for behavioral neurotoxicity focuses heavily on analyzing locomotor behavior using distance moved and population-based statistical methods (Basnet et al., 2019; G. Zhang et al., 2017). However, while the behavior repertoire of larval zebrafish is less sophisticated when compared to that of adult zebrafish and other higher-order vertebrates, they are capable of numerous distinct behaviors (Basnet et al., 2019; Kalueff et al., 2013; Mirat et al., 2013). These behaviors, such as thigmotaxis, and light avoidance cannot always be captured when using distance moved as a sole indicator of neurobehavioral toxicity in analyses of this data. Moreover, as most laboratory zebrafish populations feature significant genetic heterogeneity, individual responses to exotic toxicants cannot be expected to be homogeneous for simplistic measures such as distance moved (Balik-Meisner et al., 2018).

Improved accessibility to computing resources and application interfaces, together with recent advances in deep-learning makes it possible to analyze complex behavioral data in novel ways and predict neurodevelopmental toxicity (Arifoglu & Bouchachia, 2017; Pereira et al., 2020; Xia et al., 2018). The volume and diversity of data generated during HTS experiments, combined with the variety in toxicological response within populations, present an opportunity that is well-suited for machine learning (ML). In particular, analysis of zebrafish HTS data from five dpf larvae exposed to 1,060 unique chemicals reveals that only 8% of chemical-concentration pairs (a unique combination of chemical and concentration, e.g. 6.4 µM Nicotine) exhibit changes in distance moved (G. Zhang et al., 2017), which is alarmingly low given the known toxicity profiles of the chemical set. This challenge provides an opportunity to apply methods developed for anomaly detection from areas such as financial fraud (Awoyemi et al., 2017), medical application faults (Pachauri & Sharma, 2015), security systems intrusion (Sargolzaei et al., 2016), system faults (Warriach & Tei, 2013), and others (Fazai et al., 2019; Jaiswal & Ruskin, 2019). In anomaly detection, we learn the pattern of a normal process, and anything that does not follow this pattern is classified as an anomaly. This learning model is particularly applicable, as many HTS data sets have large amounts of control data to analyze (G. Zhang et al., 2017). One intriguing approach to achieving this is by applying an autoencoder (Feng et al., 2021; Frassek et al., 2021; Goodfellow et al., 2016; Le Borgne et al., 2022; Nicholaus et al., 2021; Ranjan et al., 2019). An autoencoder is a neural network of two modules, an encoder and a decoder (Goodfellow et al., 2016; Gupta & Singh, 2019). The encoder learns the underlying features of a process, and these features are typically in a reduced dimension. The decoder then uses this reduced dimension to recreate the original data from these underlying features.

In the present study, we trained deep autoencoder models to recognize the pattern of quintessential larval zebrafish behavior and identify abnormal behavior following developmental chemical exposure. The performance of our deep autoencoders was compared against traditional statistical methodologies, the gold standard for behavioral assessment. In addition to model development, we assessed the features driving performance through feature permutation and generated new confirmatory data to assess model reproducibility and confirm novel findings.

## Results

### Statistical classification of behavior

After classifying each of the 96-well plates by differences in the movement of controls into hyperactive, normal, or hypoactive, we compared treated vs control behavioral response to light/dark cycling in zebrafish larvae at five dpf. We identified 39 chemical-concentration combinations from ten chemicals capable of inducing a significantly different (p < 0.05) behavioral response (Supp. Table 2). Using the 30^th^ and 70^th^ percentiles, we defined 227 individual larvae as abnormal (Fig 1a). These 227 larvae formed the validation set used to test the performance of our models.

**Figure 1:**
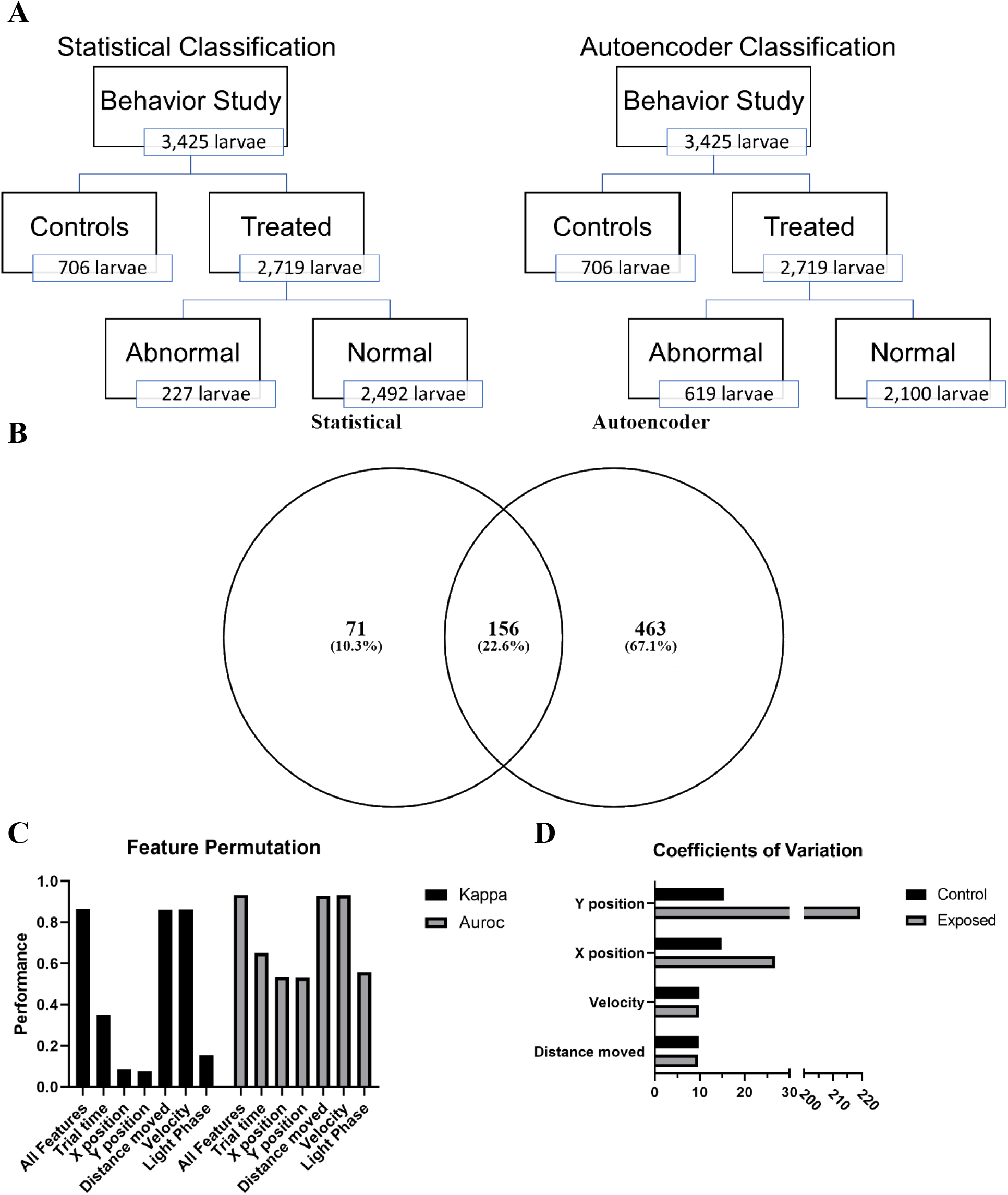
Assessment autoencoder performance. (A) Schematic representation of the differences in statistical and autoencoder based classification of behavioral response in larval zebrafish. (B) Venn diagram showing overlap between statistical and autoencoder classified abnormal zebrafish. (C) Evaluating the change in model performance when the values of a single feature are randomly shuffled. Kappa – Cohen’s Kappa statistic, AUROC - area under the receiver operating characteristic. Figure depicts means ± SEM. (D) Coefficients of variation for each of the main numerical features.

### Training performance

Autoencoder models were trained using only control data for each of the activity states (hypoactive, normal, and hyperactive) per phase of the second light cycle. This resulted in six trained models (Supp Fig 1 the training loss plots for the models). Table 1 shows the results for the six deep autoencoder models trained using control data and validated using data from zebrafish defined as abnormal using the K-S test. All the models performed well with values ranging from 0.615 – 0.867 and 0.740 – 0.922 for the Kappa and AUROC, respectively. As expected, the models consistently produced high specificity (SP) levels as this value indicated how well the models classify control data. There was greater variability in the sensitivity (SE) with the dark phase models matching or outperforming the light phase models for each activity state. Further, we observed a noteworthy trend across all models producing high positive predictive value (PPV). Overall, these results show that deep autoencoders trained using control data is capable of distinguishing between normal and abnormal larval zebrafish behavior with a high degree of accuracy.

**Table 1.**
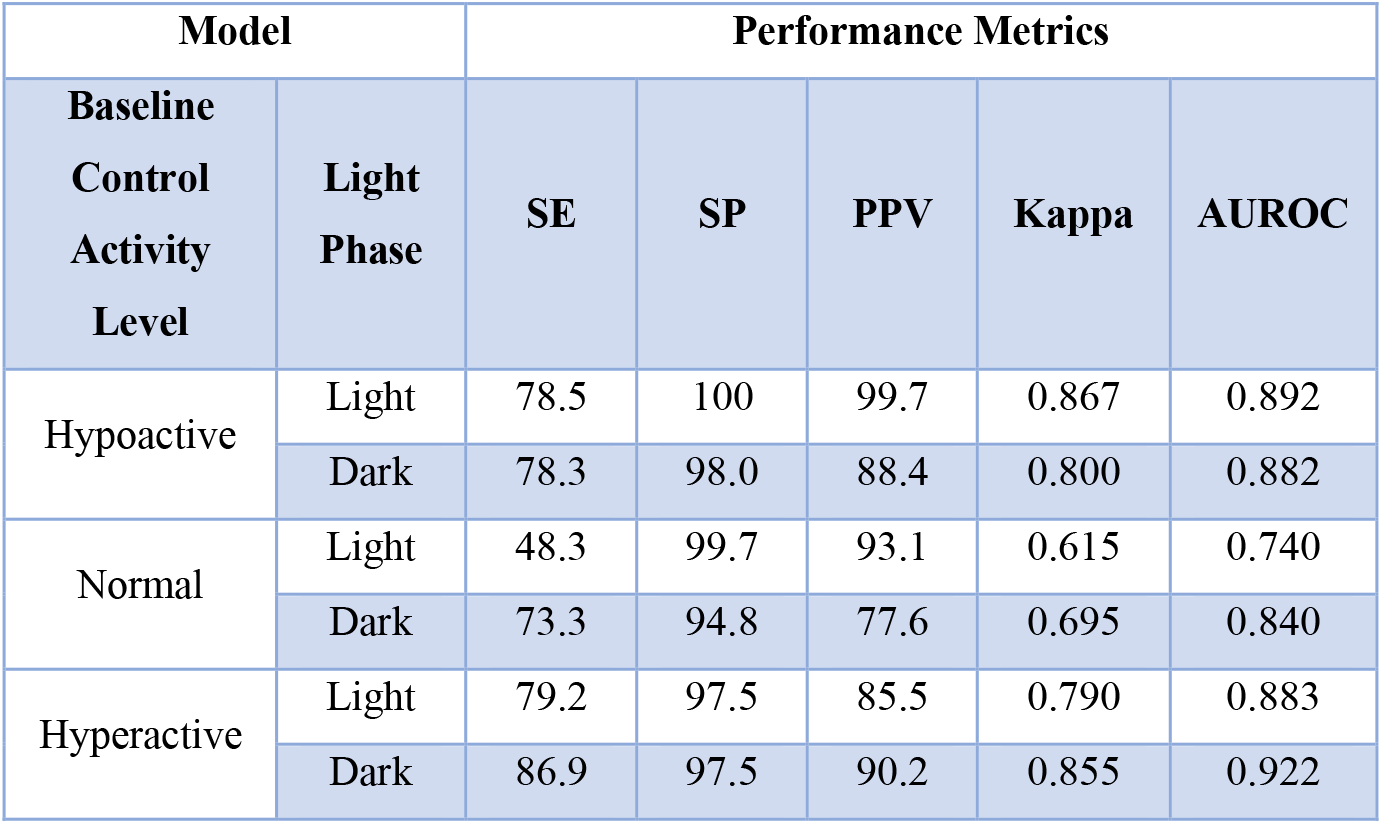
Deep autoencoder model performance in behavioral classification. Table showing performance of model trained using different activity states of the control data in both light and dark phases.

| Model |  | Performance Metrics |  |  |  |  |
| --- | --- | --- | --- | --- | --- | --- |
| Baseline Control Activity Level | Light Phase | SE | SP | PPV | Kappa | AUROC |
| Hypoactive | Light | 78.5 | 100 | 99.7 | 0.867 | 0.892 |
|  | Dark | 78.3 | 98.0 | 88.4 | 0.800 | 0.882 |
| Normal | Light | 48.3 | 99.7 | 93.1 | 0.615 | 0.740 |
|  | Dark | 73.3 | 94.8 | 77.6 | 0.695 | 0.840 |
| Hyperactive | Light | 79.2 | 97.5 | 85.5 | 0.790 | 0.883 |
|  | Dark | 86.9 | 97.5 | 90.2 | 0.855 | 0.922 |

### Evaluation of unknowns

Using the six trained models, we evaluated the 2,719 treated zebrafish larvae (Fig 1). The autoencoders correctly classified 156 of the 227 larvae that fell below or above the 30^th^ and 70^th^ percentiles, respectively. In addition, our deep autoencoders identified 463 larvae as abnormal from the 2,492 larvae defined as normal using the K-S test (Fig 1b). The majority (422) of these 619 larvae were from one of 66 chemical-concentration combinations from 13 chemicals (Table 2). The deep autoencoders successfully identified nine of the ten statistically abnormal chemicals and identified these chemicals at or below the lowest concentration shown to be statistically significant. While the deep autoencoders did not identify Perfluorodecylphosphonic acid as capable of inducing abnormal behavior, but they did identify 3-Perfluoropentyl propanoic acid (5:3), Perfluoro-n-octadecanoic acid, 8-Chloroperfluorooctylphosphonic acid, and Nonafluoropentanamide, which were missed in the statistical testing framework. These results, summarized in fig 2, show that deep autoencoders can match the performance of the K-S test and are more sensitive at detecting abnormal behavior.

**Figure 2:**
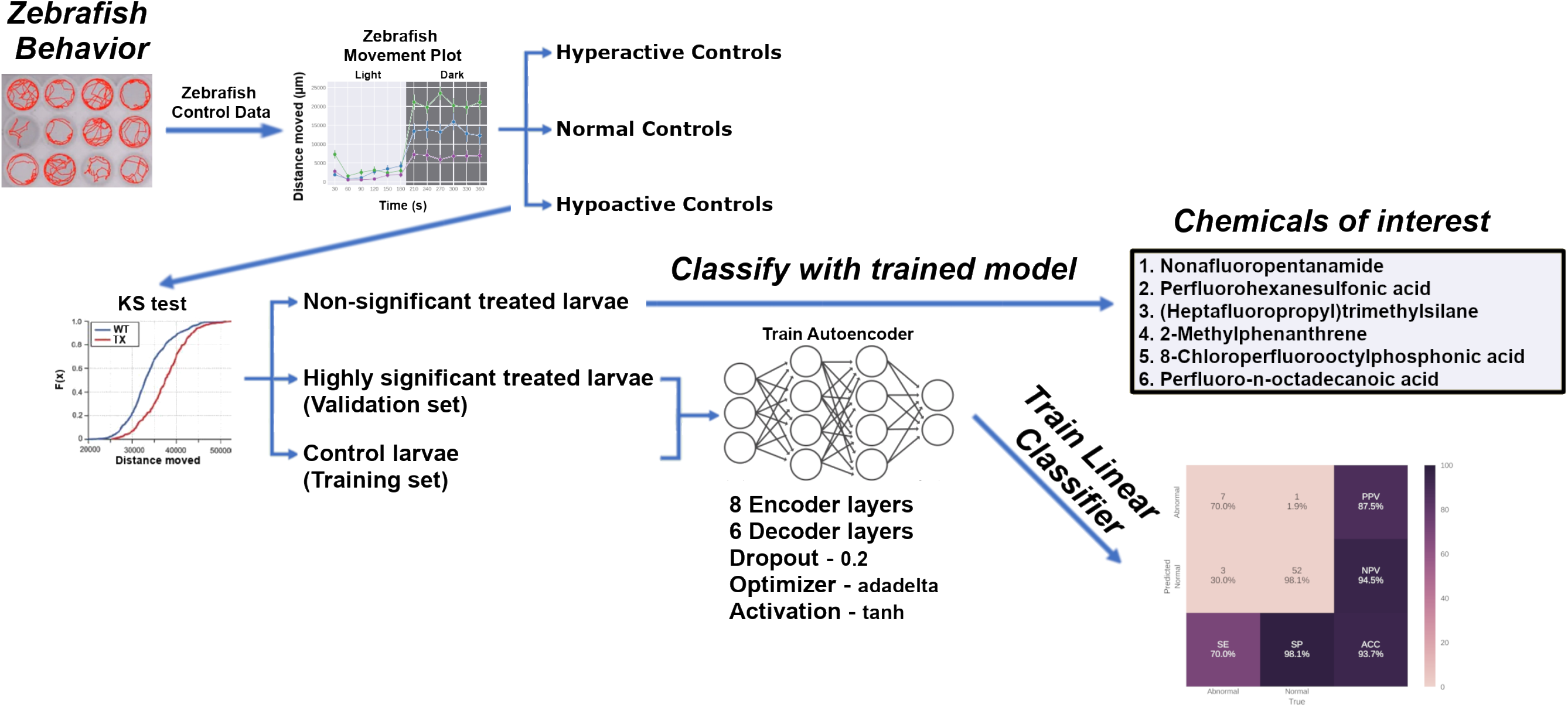
Summary of behavioral analysis pipeline and results. Utilizing our analysis pipeline produced six deep autoencoder models (three for the light phase and three for the dark phase) capable of classifying larval zebrafish behavior with high Kappa and AUROC values. The trained models were then used to classify the non-significant exposed larvae and identified Nonafluoropentanamide, Perfluorohexanesulfonic acid, (Heptafluoropropyl)trimethylsilane, 2-Methylphenanthrene, 8-Chloroperfluorooctylphosphonic acid, Perfluoro-n-octadecanoic acid, and others as capable of inducing abnormal behavior.

**Table 2.** Autoencoders identified chemicals. Table showing chemicals and concentrations flagged for displaying abnormal behavioral when evaluated using Autoencoder. Compounds that were picked up by Autoencoder, but not KS test are highlighted in red.

| CASRN | Chemical Name | Concentration (μM) |
| --- | --- | --- |
| 71751-41-2 | Abamectin | 0.1, 0.2, 0.4, 0.6 |
| 308068-56-6 | Multi-Walled Carbon Nanotube | 10, 23.2, 50, 75, 100 |
| 2531-84-2 | 2-Methylphenanthrene | 1, 2.54, 6.45, 16.4, 35, 74.8, 100 |
| 832-69-9 | 1-Methylphenanthrene | 1, 2.54, 6.45, 16.4, 35, 74.8, 100 |
| 914637-49-3 | 3-Perfluoropentyl propanoic acid (5:3) | 0.25 |
| 192-51-8 | Dibenzo[e-l]pyrene | 0.01, 0.025, 0.065, 0.164, 0.35, 0.75, 1, 2.54, 16.4, 35, 100 |
| 16517-11-6 | Perfluoro-n-octadecanoic acid | 0.25 |
| 355-46-4 | Perfluorohexanesulfonic acid | 0.015, 0.14, 0.41, 3.7, 11.1, 33.3, 66.5, 100 |
| 3834-42-2 | (Heptafluoropropyl)trimethylsilane | 0.015, 0.046, 0.41, 1.23, 11.1, 33.3 |
|  | 8-Chloroperfluorooctylphosphonic acid | 0.167 |
| 31253-34-6 | 2-Aminohexafluoropropan-2-ol | 0.015, 0.046, 0.41, 1.23, 3.7, 11.1, 33.3, 66.5, 100 |
| 13485-61-5 | Nonafluoropentanamide | 0.41, 3.7, 11.1 |
| 439-14-5 | Diazepam | 1, 3, 5, 8, 12 |

### Features driving improved autoencoder performance

To determine the features in the model that were most important in driving classification performance, we employed permutation feature importance. This technique is a model agnostic inspection technique used for any fitted estimator to determine the importance of each feature in the model. Larger the decrease in model performance (Kappa or AUROC) when a single feature value is randomly shuffled, the more important the feature. Our results, shown in fig 1c, indicate that phase, trial time, x position, and y position are the largest drivers of model performance, while distance moved and velocity contribute very little. Coefficients of variation show greater variability in the x and y positional data between control and exposed groups compared to either velocity or distance moved (fig 1d). This trend is consistent irrespective of the larval activity state (hypoactive, normal activity, or hyperactive) relative to their respective controls (Fig 3).

**Figure 3:**
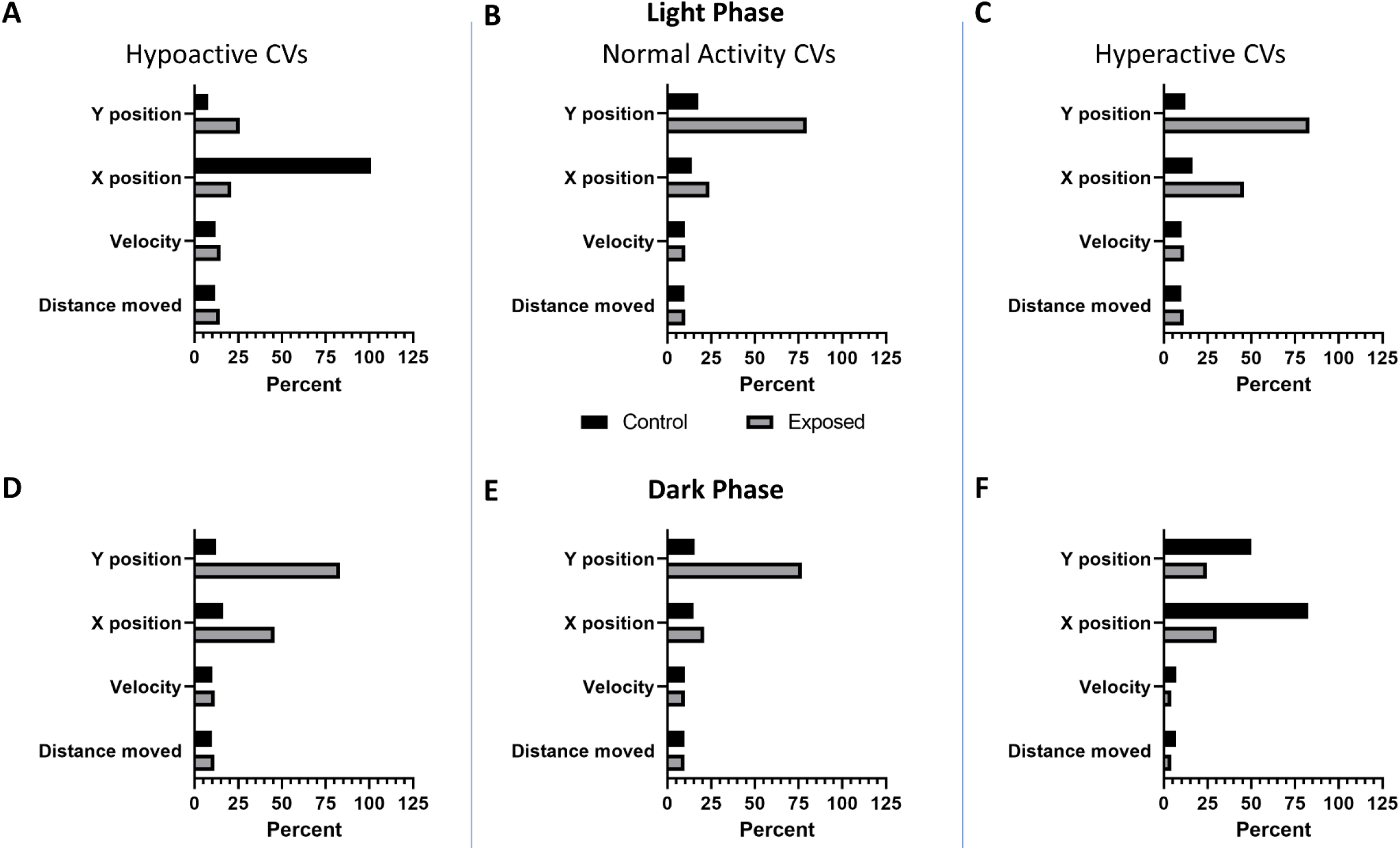
Coefficients of variation per larval activity state. Coefficients of variation (CVs) for each of the main numerical features (A – C) in the light (D – F) and in the dark. Columns show CVs of larval zebrafish significantly (p < 0.05) (A, D) hypoactive, (B, E) normal activity, or (C, F) hyperactive relative to their respective controls.

### Experimental confirmation of autoencoder findings

To provide an unbiased evaluation of the final model fits, we generated new data using 2-Methylphenanthrene, and Nonafluoropentanamide. The data collected confirmed that our models accurately classified all controls as normal while detecting similar levels of abnormal behavior response across the concentration range (Fig 4). These results show that the trained model is capable of producing similar results across experimental replicates.

**Figure 4:**
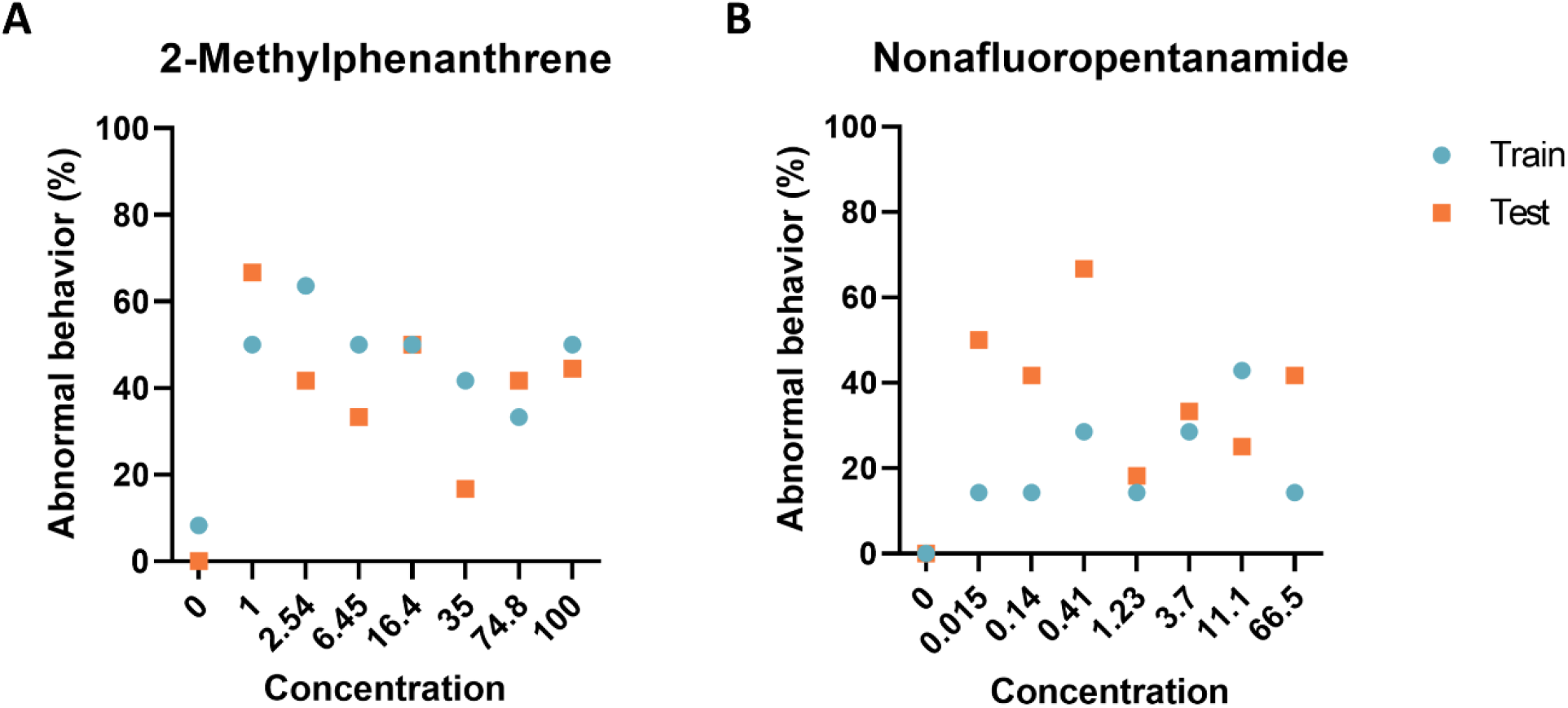
Experimental model evaluation. Comparison of the performance of deep autoencoder models between the training set and two chemicals identified by the models to elicit abnormal larval zebrafish behavior. Percent of larval zebrafish classified as abnormal based on their behavioral response to developmental exposure to (A) 2-Methylphenanthrene and (B) Nonafluoropentanamide

## Discussion

Statistical analysis identified 39 chemical-concentration combinations from ten chemicals capable of inducing a significantly different (p < 0.05) behavioral response. Utilizing the 227 abnormal individuals identified by the statistical test as our validation set, we trained six deep autoencoder models using control data for each of the activity states (hypoactive, normal, and hyperactive). All of the resulting models performed well with values ranging from 0.615 – 0.867 and 0.740 – 0.922 for the Kappa and AUROC, respectively. All models achieved SP values above 94.8% and PPV values above 77.6% while SE values for all dark phase models outperformed the light phase models for each activity state (Table 1). Assessment of permutation feature importance indicates that phase, trial time, x-position, and y-position are the largest drivers of model performance (fig 1c). The calculated coefficients of variation shed some light on this surprising finding (fig 1d). They show that variation in the x and y positional data is greater than observed for velocity or distance moved between control and exposed groups. These differences in variation likely make it easier for the models to distinguish between treated and exposed groups.

When we examined exposed larvae defined as normal using the K-S test (Fig 1), our deep autoencoders identified 66 chemical-concentration combinations from 12 chemicals (Table 2) with Perfluoro-n-octadecanoic acid, 8-Chloroperfluorooctylphosphonic acid, and Nonafluoropentanamide only identified by our autoencoders. These results show that a deep autoencoder-based model can classify larval zebrafish behavior as normal or abnormal with very good efficacy and often identified abnormal behaviors at lower concentrations than current statistical methods. Further, the models identified three novel chemicals, Perfluoro-n-octadecanoic acid, 8-Chloroperfluorooctylphosphonic acid, and Nonafluoropentanamide as capable of inducing abnormal behavior (Fig 3).

Recognition and categorization of swimming patterns in larvae is a challenging task and a number of approaches have been used. These can range from subjective analysis based on experienced observations (Fero et al., 2011; Kalueff et al., 2013, p. 0) or more recently through the application of unsupervised ML (Budick & O’Malley, 2000; Burgess & Granato, 2007a, 2007b, 2007c; Kimmel et al., 1974; Mirat et al., 2013; H. Zhang et al., 2013). These studies have focused on the analysis and categorization of behavioral patterns in wild-type strains (Burgess & Granato, 2007c; H. Zhang et al., 2013), mutant strains (Burgess & Granato, 2007b; Mirat et al., 2013), or larvae exposed to neuroactive chemicals (Mirat et al., 2013) but do not classify behavior as normal or abnormal. In addition, these unsupervised approaches have utilized highspeed camera systems which are medium to low throughput and have limited potential in the screening of tens of thousands of chemicals for behavioral effects. As introduced above, classification of behavior is one of the primary goals of toxicological screening and tends to result in highly imbalanced datasets and lend themselves to anomaly detection methodologies. While these methods are common in manufacturing (Fan et al., 2018; Fazai et al., 2019; Jaiswal & Ruskin, 2019; Nicholaus et al., 2021), information systems (Pachauri & Sharma, 2015; Warriach & Tei, 2013), security systems (Feng et al., 2021; Sargolzaei et al., 2016), and financial fraud (Awoyemi et al., 2017) they have only very recently been applied to biological data (Frassek et al., 2021; Homayouni et al., 2021; Nwokedi et al., 2021). To the best of our knowledge, this is the first study to explicitly develop a deep autoencoder model for anomaly detection in toxicological behavior studies.

Overall, our results show that a deep autoencoder utilizing raw behavioral tracking data from five dpf zebrafish larvae can accurately distinguish between normal and abnormal behavior. We show that these results are reproducible and allow for the identification of new compounds capable of eliciting abnormal behavior. Further, our models were able to identify abnormal behavior following chemical exposure at lower concentrations than with traditional statistical tests. Our approach accounts for the high degree of behavioral variability associated with the genetic diversity found within a highly outbred population typical of zebrafish studies, thereby making it extensible to use across labs. Looking to the future, neurodevelopmentally active chemicals identified using our deep autoencoder models may represent heretofore undetectable signals of subtle differences in individual responses, suggesting chemicals that should be investigated further as eliciting differential population responses (i.e. interindividual susceptibility differences).

These findings will facilitate the application of behavioral characterization methods discussed above, such as Zebrazoom (Mirat et al., 2013), using highspeed cameras to identify the behavioral traits most perturbed by the chemical exposure and allow for more mechanistic discovery. One of the key innovations presented in this study is leveraging vast amounts of control data generated as part of any high-throughput screening (HTS) – setting the stage for predictive toxicological applications and safety assessments for the enormous backlog of as-yet untested chemicals.

## Materials and methods

This section describes the autoencoder models utilizing a semi-supervised ML algorithm and logistic regression (LR) to discriminate between normal and abnormal behavior in chemically exposed five dpf zebrafish. An overview of our approach is shown in Fig 3. Briefly, we created and trained six autoencoder models for each phase of the assay; namely, hyperactive, normal, and hypoactive depending on the control movement in the light or dark phases of the assay. Finally, treated plates were tested on one of these, depending on which category, its controls fell under. We used experimental data collected on a large and diverse compound set of 30 chemicals including an insecticide, nanomaterial, perfluorinated chemicals, and aromatic pollutants at a range of concentrations (133 chemical-concentration pairs) to assess the neurotoxic effects of these chemicals following developmental exposure (Supp. Table 1).

### Zebrafish husbandry

Tropical 5D wild-type zebrafish were housed at Oregon State University’s Sinnhuber Aquatic Research Laboratory (SARL, Corvallis, OR) in densities of 1000 fish per 100-gallon tank according to the Institutional Animal Care and Use Committee protocols (Barton et al., 2016). Fish were maintained at 28 °C on a 14:10 h light/dark cycle in recirculating filtered water, supplemented with Instant Ocean salts. Adult, larval and juvenile fish were fed with size-appropriate GEMMA Micro food 2–3 times a day (Skretting). Spawning funnels were placed in the tanks the night prior, and the following morning, embryos were collected and staged (Kimmel et al., 1995; Westerfield, 2007). Embryos were maintained in embryo medium (EM) in an incubator at 28 °C until further processing. EM consisted of 15 mM NaCl, 0.5 mM KCl, 1 mM MgSO_4_, 0.15 mM KH_2_PO_4_, 0.05 mM Na_2_HPO_4_, and 0.7 mM NaHCO_3_ (Westerfield, 2007).

### Developmental chemical exposure

The empirical data used to develop our model were gathered as described in Truong et al. and Noyes et al.(Noyes et al., 2015; Truong et al., 2014, 2022). The experimental design consisted of the 30 unique chemicals tested (Supp Table 1) with at least 7 replicates (an individual embryo in singular wells of a 96-well plate) at each concentration for each chemical.

### Developmental toxicity assessments

#### Mortality and morphology

At 24 hours post-fertilization (hpf), embryos were screened for mortality and 2 developmental endpoints. At 120 hpf, mortality and incidence of abnormality in 9 morphology endpoints were evaluated as binary outcomes. Any individuals identified with a physical abnormality were excluded from any behavioral analysis as these abnormalities might confound the results.

#### Photomotor responses

The larval photomotor response (LPR) assay was conducted at 120 hpf when the 96-well plates of larvae were placed into a Zebrabox (Viewpoint LifeSciences) and larval movement was recorded. The recorded videos were then tracked with Ethovision XT v.11 analysis software for 24 min across 3 cycles of 3 min light: 3 min dark. The trial time(s), x-position, y-position, distance moved (µm), and velocity (mm/s) by each larva in the 2nd light/dark cycle were the features used for behavioral assessment (Supp Fig 2). The 2^nd^ light/dark cycle was chosen as it exhibited less noise than the 1^st^ cycle and was less influenced by any learning that might have occurred in the 3^rd^ cycle. For all assessments, data were collected from embryos exposed to nominal concentrations of chemical and uploaded under a unique well-plate identifier into a custom LIMS (Zebrafish Acquisition and Analysis Program [ZAAP]) – a MySQL database and analyzed using custom R scripts that were executed in the LIMS background (Truong et al., 2016).

### Data preprocessing and statistical analysis pipeline

#### Preprocessing

All data processing, statistical analysis and ML were implemented in Python using the open source libraries Tensorflow (Martín Abadi et al., 2015), Keras (U.S. Environmental Protection Agency, 2021), Scikit-learn (Pedregosa et al., 2011), Pandas (McKinney, 2010), and Numpy (Harris et al., 2020) within a purpose build Singularity container environment (Sylabs.io, 2019). The x-position and y-position data was standardized relative to the center of each well and forward filled if datapoints were missing. Outliers were normalized to the maximum likely distance a zebrafish larva could move in 1/25^th^ of a second. Considering that the average length of a 5 dpf larval zebrafish is 3.9 mm and can move about 2.5 times it’s body length during a startle response (120 frames at 1000 frames/second) the threshold for distance moved in our system was set at 3.25 mm per frame (Burgess & Granato, 2007b; *ZFIN Zebrafish Developmental Stages*, n.d.). This resulted in 5,445 of the 30,825,000 frames being normalized.

#### Statistical analysis

Interexperimental zebrafish larval response to light/dark cycling is highly variable (Supp Fig 2). Therefore, a two sample Kolmogorov–Smirnov test (K-S test) was used to compare mean of controls from individual 96-well plates to mean control movement across all plates. The K-S test is a non-parametric two-sided test and no adjustments were made for normality or multiple comparisions. Controls from individual plates with statistically significant (p < 0.01) differences in movement compared to the average of all controls were grouped together as hyperactive, normal, or hypoactive. Following grouping the K-S test was used to compare each chemical-concentration combination with their respective same plate control (p < 0.05). Individuals in the 30^th^ and 70^th^ percentiles of each chemical-concentration combination were defined as abnormal.

### Autoencoder architecture

Deep autoencoders were developed using zebrafish control data to distinguish between normal and abnormal zebrafish behavior. The model was trained on a Dell R740 containing two Intel Xeon processors with 18 cores per processor, 512 GB RAM, and a Tesla-V100-PCIE (31.7 GB). The autoencoders consisted of an input and output layer of fixed-size based on the size of a single phase (25 frames per 180s) of the second light cycle (4500 frames by 5 features). The encoder network was composed of eight fully connected hidden layers using a normal kernel initialization, tanh activation, a dropout value of 0.2, L1 and L2 regularization values of 1e^-05^, and an adadelta optimizer. The size of each hidden layer was reduced by increasing multiples of 15 and resulted in a compressed representation (bottleneck) size of 250. The decoder network was composed of six fully connected hidden layers using tanh activation, and a dropout value of 0.2. All hidden layers used an adadelta optimizer (learning_rate=0.001, rho=0.95, and epsilon=1e-07) and mean squared error for the loss function (He et al., 2015; Osl et al., 2012; Ramachandran et al., 2017). For each model, we optimized the hyperparameters (i.e., the number of hidden layers, the number of nodes in the layers, loss functions, optimizers, regularization rates, and dropout rates) by grid search technique trained on all control data over 500 epochs using Cohens Kappa statistic as the objective metric. The final encoder models were trained over the course of 125000 epochs. The resulting compressed representation was used as input into a logistic regression layer trained using a 100 fold cross-validation with each fold consisting of 4000 epochs using a limited-memory BFGS solver. The code and sample training data that implements the models are available at GitHub [https://github.com/Tanguay-Lab/Manuscripts/tree/main/Green_et_al_(2023)_Manuscript]. A complete dataset is available apon request.

### Network performance and evaluation

The data showed strong normal vs abnormal class imbalance (Fig 1). Classifiers may be biased towards the major class (normal) and therefore, show poor performance accuracy for the minor class (abnormal) (Lemaître et al., 2017). Normal vs abnormal classification accuracy was evaluated using a confusion matrix, Cohen’s Kappa statistic, and area under the receiver operating characteristic (AUROC) as Kappa and AUROC measure model accuracy, while compensating for simple chance (Ben-David, 2008). The primary metrics we used from the confusion matrix included sensitivity (SE), specificity (SP), and positive predictive value (PPV) as these parameters give us the true positive rate, true negative rate, and the proportion of true positives amongst all positive calls (Parikh et al., 2008; Pearson, 1904; Townsend, 1971). Chemical-concentration combinations were defined as abnormal if the autoencoders identified more individual as abnormal in the exposed than their respective controls and at least 25% of the individuals were abnormal. Permutation feature importance was used to evaluate which features are the most important for model performance. In brief, one feature (variable) is shuffled randomly and all features are fed into the model the resulting Kappa and AUROC values are calculated. This is repeated 1000 times per feature and average Kappa and AUROC are calculated across each shuffle (Breiman, 2001). To determine why one feature might be more important than another a coefficient of variation was calculated for each of the features in the control and exposed groups (variation() in the Scipy package).

## Acknowledgments

This research was supported by the National Institutes of Health, through the National Institute of Environmental and Health Sciences (P30 ES030287, R56 ES030007, P30 ES025128) and the National Cancer Institute (R01 CA161608). We would like to thank the staff at Sinnhuber Aquatic Research Laboratory, and John Lam for his contribution to reprocessing videos.

